# Multimodal Detection of Dopamine by Sniffer Cells Expressing Genetically Encoded Fluorescence Sensors

**DOI:** 10.1101/2021.09.16.460471

**Authors:** Carmen Klein Herenbrink, Jonatan Fullerton Støier, William Dalseg Reith, Abeer Dagra, Miguel Alejandro Cuadrado Gregorek, Yulong Li, Lin Tian, Ulrik Gether, Freja Herborg

## Abstract

Dopamine serves an important role in supporting both locomotor control and higher brain functions such as motivation and learning. Dopaminergic dysfunction is implicated in an equally multidimensional spectrum of neurological and neuropsychiatric diseases. Extracellular dopamine levels are known to be tightly controlled by presynaptic dopamine transporters (DAT), which is also a main target of psychostimulants. Still, detailed data on dopamine dynamics in space and time is needed to fully understand how dopamine signals are encoded and translated into cellular and behavioral responses, and to uncover the pathological effects of dopamine-related diseases. The recently developed genetically encoded fluorescent dopamine sensors enable unprecedented monitoring of dopamine dynamics and have changed the field of *in vivo* dopamine recording. However, the potential of these sensors to be used for *in vitro* and *ex vivo* assays remains unexplored. Here, we demonstrate a generalizable blueprint for making “sniffer” dopamine cells for multimodal detection of dopamine *in vitro* and *ex vivo*. We generated sniffer cell lines with inducible expression of six different dopamine sensors and performed a head-to-head comparison of sensor properties to guide users in sensor selection. In proof-of-principle experiments, we show how the sniffer cells can be applied to measure release of endogenous dopamine from cultured neurons and striatal slices, and for determining total dopamine content in striatal tissue. Furthermore, we use the sniffer cells to quantify DAT-mediated dopamine uptake, and AMPH-induced and constitutive dopamine efflux as a radiotracer free, high-throughput alternative to electrochemical- and radiotracer-based assays. Importantly, the sniffer cells framework can readily be applied to other transmitter systems for which the list of genetically encoded fluorescent sensors is rapidly growing.

## Introduction

Dopamine (DA) serves as a neuromodulator in the brain where it is critically involved in locomotor control and higher brain functions such as motivation and reward-related learning. In line with these functions, decades of research have implicated disturbances in dopaminergic neurotransmission in both movement disorders and mental illnesses ^1^. The dopaminergic circuit of the brain is also a major target for several therapeutics used in the treatment of these movement and mental disorders, and for drugs of abuse ^2, 3, 4, 5^. Still, we only have a limited understanding of the nature and progression of DA dysfunction in diseased states. In addition, the mechanisms through which DA exerts its short- and long-term effects on emotional states and behavior remain unclear. Sensitive methods to study DA neurotransmission in cell cultures, tissue preparations, and living organisms are necessary to gain mechanistic insights into DA signaling in health and disease states, and for the development of effective therapeutics that target dopaminergic circuits.

Important advances in the field of DA detection have recently been achieved with the development of G protein-coupled receptor (GPCR)-based sensors that directly couple the presence of DA with an increase in fluorescence. These sensors allow the interrogation of extracellular DA levels with unprecedented spatiotemporal resolution using optical measurements of fluorescence intensity ^6, 7^. Several studies have already demonstrated the powerful application of viral expression of such sensors for studying DA dynamics in live animals and in brain tissue circuits using e.g. fiber photometry and microscopy techniques ^6, 7, 8, 9, 10, 11, 12^. Two families of GPCR-based DA sensors are currently available, the dLight and GRAB_DA_ family, which are based on the DA D_1_ receptor (D1R) and DA D_2_ receptor (D2R), respectively. While both of these sensor families are based on the coupling of inert DA receptors to a conformational sensitive circularly permuted GFP molecule, they encompass distinct properties ^6, 7, 13, 14^. Overall, the sensors fulfill a number of attractive features such as high molecular specificity and affinities similar to that of endogenous DA receptors, large dynamic ranges, and a single-fluorophore protein design that allows for viral delivery and cell specific expression ^6, 7, 13, 14^. However, the sensors have different intrinsic properties that should be carefully considered in context of experimental conditions and the research question, but a direct side-by-side comparison of dopamine sensors is currently lacking to guide neuroscientists in choosing the most suitable sensor. In addition, the potential use of the growing list of GPCR-based sensors for development of assays for *in vitro* and *ex vivo* DA recordings is largely unexplored.

Here, we present an easy to use, inexpensive, and scalable framework for applying GPCR-based DA sensors for multimodal *in vitro* and *ex vivo* measurements of DA. We establish six different DA “sniffer” cell lines with inducible expression of three dLight and three GRAB_DA_ sensors and carry out a head-to-head comparison of sensor properties under identical experimental conditions. We perform proof-of-principle experiments showing how such sniffer cells can readily be applied to record release of endogenous DA from cultured neurons and striatal slices, and to determine total DA content in striatal tissue. Moreover, we demonstrate that the sniffer cells also enable measurements of DA transporter (DAT) activity, such as DA uptake and efflux, allowing for a radiotracer free, high-throughput alternative to electrochemical- and radiotracer-based assays. Importantly, this framework for versatile usage of DA sniffer cells can easily be applied to other transmitter systems for which the palette of genetically encoded single-fluorescent protein sensors is continuously expanding. Because of the ease of use, low costs, and virus- and radioactivity-free properties, fluorescent sensor-expressing sniffer cells have great potential for becoming a general tool for studying transmitter levels in culture systems and tissue preparations.

## Results and Discussion

### Development and Characterization of DA Sniffer Cell Lines

With the novel development of GPCR-based DA sensors, we wanted to expand the toolbox for DA detection *in vitro* and *ex vivo* with a virus and radiotracer-free method that allows for DA detection across multiple assay and sample formats using commonly available plate readers and fluorescence microscopes. To do this, we generated DA “sniffer” cells by stable transfection of Flp-In T-REx 293 cells with DA sensors of either the dLight or the GRAB_DA_ sensor family. Six DA sensors were selected for this study: dLight1.1, dLight1.2, dLight1.3a, GRAB_DA1M_, GRAB_DA1H_, and GRAB_DA2M_. Of note, as the Flp-In system was utilized for the generation of the sniffer cell lines, all sensors were inserted at the same specific genomic location ^15^.

We first validated the sniffer cell lines using fluorescence microscopy to ensure tetracycline-induced expression of the sensors and confirm DA sensitivity. Indeed, upon treatment with tetracycline, all cell lines expressed their respective sensor, and all displayed an increase in fluorescence upon incubation with 10µM DA (**Figure 1A**). We then carried out a head-to head comparison of the six different DA sniffer cell lines to derive key sensor properties under identical experimental conditions. The most important sensor properties one needs to consider are the dynamic range, sensor sensitivity, and kinetic parameters, which need to be compatible with the expected DA concentrations and fluctuations of the model system.

**Figure 1.**
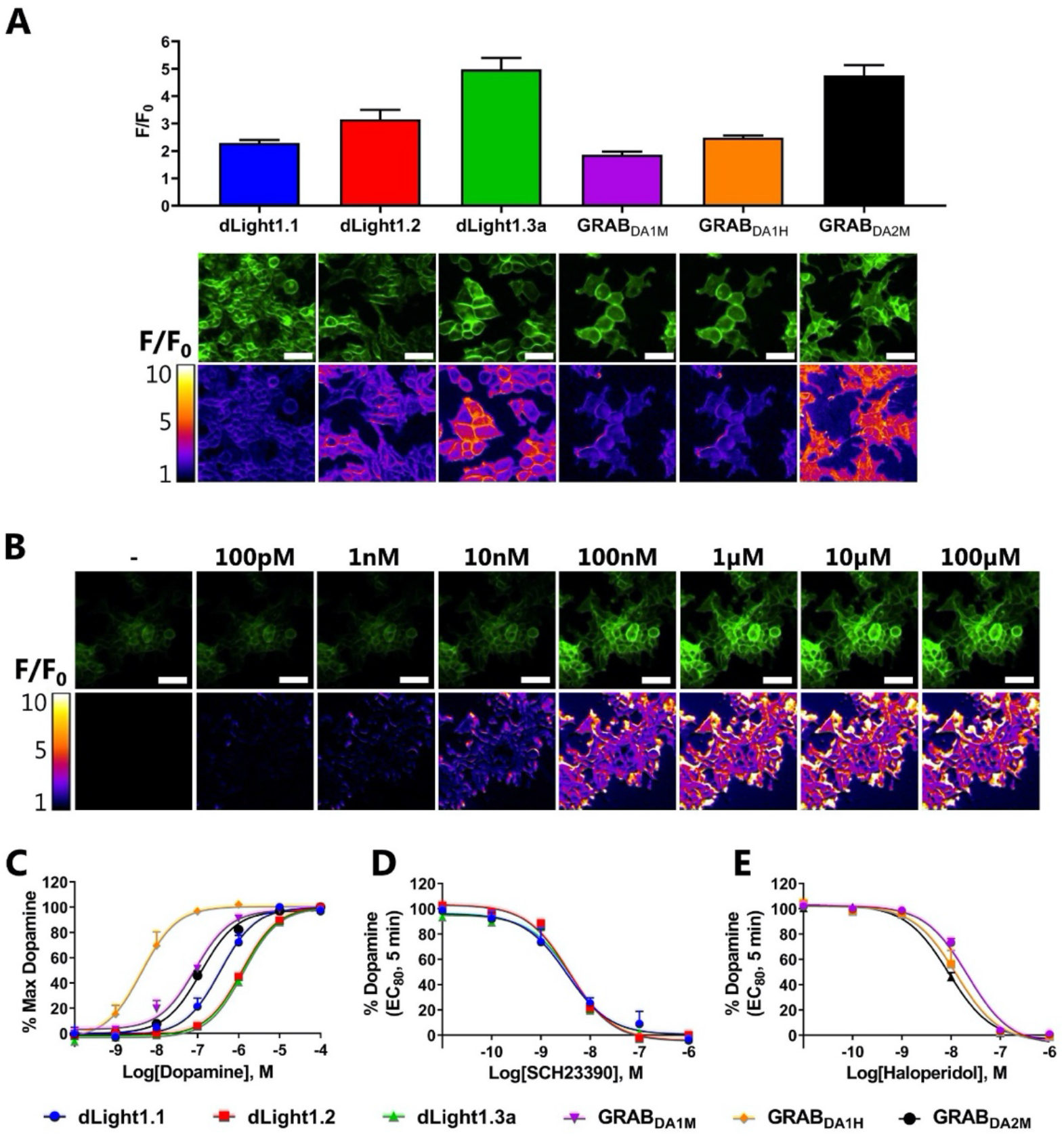
Characterization of the Dynamic and Detection Range of DA Sniffer Cell Lines. **A)** Visualization and quantification of the fluorescent change in Flp-In T-REx-293 cells expressing one of six different DA sensors. Cells were stimulated with 10µM DA while imaged on an epifluorescence microscope (N=3, mean ± SEM). Representative images of the raw fluorescent signal (top) are shown along with pseudo colored representations of the fold change above baseline (bottom). Scale bars are 50µM. **B)** Representative images of GRAB_DA2M_ sniffer cells stimulated with increasing doses of DA show a dose-dependent increase in fluorescence (N=3). Similar dose-response images of the remaining sniffer cell lines can be found in **Supplementary Figure 1**. Scale bars are 50µM. **C)** DA dose-response curves for the six DA sniffer cell lines recorded after 5 min stimulation at 37°C with indicated DA concentrations using a fluorescence plate reader. **D, E)** The antagonists SCH23390 (**D**) and haloperidol (**E**) dose-dependently blocked the DA-induced increase in fluorescence in sniffer cells expressing the DA D_1_ and D_2_ receptor-derived sensors, respectively, as detected by a fluorescence plate reader. The values are expressed as mean ± SEM from three independent experiments.

To determine the dynamic range, we applied fluorescence microscopy to measure the change in fluorescence (F/F_0_) following application of 10µM DA to the sniffer cells (**Figure 1A, Table 1**). The greatest dynamic range was observed for the dLight1.3a (F/F_0_ = 4.98 ± 0.24) and GRAB_DA2M_ (F/F_0_ = 4.77 ± 0.22) sensors, while dLight1.1 (F/F_0_ = 2.29 ± 0.06) and GRAB_DA1M_ (F/F_0_ = 1.86 ± 0.07) sensors showed the lowest dynamic range. dLight1.2 and GRAB_DA1H_ had a fluorescent change (F/F_0_) of 3.16 ± 0.20 and 2.49 ± 0.04, respectively. The dynamic range for the D2R-derived sensors (GRAB_DA_ family) was similar to previous studies ^7, 16^. We also observed comparable, albeit slightly smaller, dynamic ranges in the sniffer cells expressing the D1R-derived sensors (dLight family) as compared to what was previously reported ^6^. The small changes in dynamic ranges can likely be explained by a difference in background fluorescence (F_0_) as a result of differential experimental set-up and/or differences in expression levels of the sensors. Importantly, however, the relative difference in F/F_0_ between the three dLight sensors appears to be consistent between the studies, with for instance dLight1.3a showing a two-fold greater F/F_0_ than dLight1.1.

**Table 1.** DA sensor Parameters.

| Sensor | pEC<sub>50</sub> ± SEM (nM)<sup>a</sup> | Detection Range | Dynamic Range | Kinetic Parameters |  |
| --- | --- | --- | --- | --- | --- |
|  |  | 10% - 90% (nM)<sup>a</sup> | F/F<sub>0</sub> ± SEM<sup>b</sup> | k<sub>on</sub> (M<sup>-1</sup> min<sup>-1</sup>)<sup>a</sup> | k<sub>off</sub> (min<sup>-1</sup>)<sup>a</sup> |
| dLight1.1 | 6.45 ± 0.12 (350) | 40 – 3100 | 2.29 ± 0.06 | 1.84 ± 0.76 x 10<sup>8</sup> | 120 ± 17 |
| dLight1.2 | 5.92 ± 0.04 (1200) | 140 – 10000 | 3.16 ± 0.20 | 7.50 ± 0.59 x 10<sup>7</sup> | 130 ± 17 |
| dLight1.3a | 5.87 ± 0.06 (1300) | 130 – 14000 | 4.98 ± 0.24 | 6.42 ± 0.72 x 10<sup>7</sup> | 150 ± 4.8 |
| GRAB<sub>DA1M</sub> | 7.12 ± 0.04 (75) | 4.0 – 1400 | 1.86 ± 0.07 | 6.60 ± 1.66 x 10<sup>8</sup> | 80 ± 6.0 |
| GRAB<sub>DA1H</sub> | 8.34 ± 0.19 (4.6) | 0.78 – 27 | 2.49 ± 0.04 | ND | 9.2 ± 0.42 |
| GRAB<sub>DA2M</sub> | 6.90 ± 0.07 (130) | 8.6 – 1800 | 4.77 ± 0.22 | 3.14 ± 0.38 x 10<sup>8</sup> | 45 ± 4.3 |

<sup>a</sup>Determined with a fluorescence plate reader
<sup>b</sup>Determined with a fluorescence microscope
<sup>ND</sup> Not determine due to ligand depletion

Next, we characterized the sensors’ DA sensitivity by exposing the sniffer cells to increasing DA concentrations. The fluorescent change was detected with both a fluorescence microscope and plate reader to determine the detection range (**Figure 1B-C, Supplementary Figure 1**). As is evident from the concentration-response curves in **Figure 1C**, the different sensors can detect DA at a wide range of concentrations. The sniffer cells expressing the D1R-derived sensors have a detection range of 40 nM to 14 µM, with the dLight1.1 sniffer cells being the most sensitive. The D2R-derived sensors, on the other hand, were more sensitive to DA, which is consistent with the D2R having a higher DA affinity than the D1R. The GRAB_DA1M_- and GRAB_DA2M_ sniffer cells had a detection range of 4 nM to 1.8 µM DA, whereas the GRAB_DA1H_ sniffer cells were able detected DA levels as low as 1 nM (**Table 1**). Importantly, we confirmed that the increases in fluorescence were mediated via the sensors, as their responses were blocked by selective DA receptor antagonists (**Figure 1D-E**). Overall, the observed detection ranges were similar between plate reader and microscope experiments and in line with previous studies ^6, 7, 13^.

Activation and deactivation kinetics of a sensor are pivotal parameters to consider for experiments where detection of DA fluctuations at high temporal resolution is important. For instance, temporal resolution is critical to capture the rapid dynamics of DA release and clearance from dopaminergic neurons. To gain greater insight into the kinetics of the sensors utilized in this study, we determined the on (*k*_on_) and off (*k*_off_) activation rates by DA at the various sensors. To do so, we stimulated the sniffer cells with two relatively low concentrations of DA (to ensure that the kinetics are activation and not diffusion driven) and measured the change in fluorescence over time (**Table 1** and **Supplementary Figure 2A-E**). It should be noted that previous studies have already determined time (τ) or half-life (t_1/2_) constants for some of the sensors utilized in our study. However, as the on-rate is dependent on the DA concentration, such values cannot be compared between sensors. As shown in **Table 1**, we found that the D1R-based dLight sensors displayed slower on-rates (*k*_on_) than the D2R-based GRAB_DA1M_ and GRAB_DA2M_ sensors. The off-rates (*k*_off_), however, were faster for the dLight sensors than the GRAB_DA_ sensors (**Table 1**). Through deep brain imaging, Patriarchi *et al*. (2018) previously determined *in vivo* that the dLight1.1 and dLight1.2 sensors display decay half-life constants of 100 ms and 90 ms, respectively ^6^. We derived off-rates (*k*_off_) of 120 ± 17 min^−1^ and 130 ± 17 min^−1^ for dLight1.1 and dLight1.2, respectively, which equal to decay half-life constants (t_1/2_) of 340 ms and 324 ms (t_1/2_ = ln2 / *k*_off_), respectively. The slower off-rate observed in our study may arise from the differential experimental conditions under which the parameters were obtained (HEK293 cells activated by DA addition versus optogenetic activation of dopaminergic neurons in live animals). The off-rates of GRAB_DA1M_ (*k*_off_ = 80 ± 6.0 min^−1^) and GRAB_DA2M_ (*k*_off_ = 45 ± 4.3min^−1^) were similar to what has previously been reported (85 min^−1^ and 46 min^−1^, respectively (*k* = τ^−1^)) ^7, 16^. Unfortunately, we were not able to obtain accurate on- and off-rates for GRAB_DA1H_ likely due to ligand depletion caused by the high sensitivity and expression of the sensor in combination with a small assay volume. To overcome ligand depletion, we determined the dissociation kinetics in the presence of a high concentration of the D2R antagonist haloperidol instead (**Supplementary Figure 2F**). We obtained an off-rate of 9.2 ± 0.42 min^−1^ for the GRAB_DA1H_ sensor which was slower than what previously has been reported (24 min^−1^ (*k* = τ^−1^)) ^7, 16^.

In summary, our head-to-head comparison of DA sensors on the detection range, dynamic range, and kinetic parameters should serve as a guideline for users to select the most appropriate DA sensor to use in their specific experiments. In assays with poor signal-to-noise ratio, the large dynamic range of dLight1.3a and GRAB_DA2M_ is necessary to accurately measure extracellular DA levels. On the other hand, high DA sensitivity is an attractive property for assays that require detection of low DA concentrations. For example, if one needs to measure very low levels of extracellular DA (< 4nM), GRAB_DA1H_ is one of the few sensors currently available that has the required DA sensitivity. However, the increased DA sensitivity is associated with a slower off-rate, which reduces the temporal detection accuracy due to integration of temporally close release events. Thus, for *in vivo* and *ex vivo* experiments where it is paramount to capture the rapid firing events of DA neurons and where DA levels are sufficiently high, the fast off-rate kinetics of the dLight sensors are favored. Another important consideration related to the sensors’ kinetic properties is their potential buffering effect, where DA availability to endogenous receptors is altered by the sensor expression. High expression of sensors, particularly with slow off-rates, may buffer a significant fraction of extracellular DA and blunt fast changes in dopamine levels, which could influence cellular or behavioral responses. Collectively, the distinct properties of each DA sensor should be carefully considered and matched to the technical setup and biological conditions.

### DA Sniffer Cells: A Novel Tool for the Study of DAT Pharmacology

DAT plays an important role in DA homeostasis as it rapidly clears DA from the extracellular space to the cytoplasm for subsequent storage and release. DAT is also the primary target for both illicit substances (e.g., psychostimulants such as cocaine and methamphetamine) and therapeutic agents such as amphetamine (AMPH) and methylphenidate used for treatment of ADHD ^17^. We wanted to assess if the DA sniffer cells could be used as a novel tool to study DAT function. For this, we transiently transfected GRAB_DA1M_ sniffer cells with human DAT (hDAT). We rationalized that DA taken up into the cells by hDAT would no longer be available extracellularly to activate GRAB_DA1M_. Thus, exposure to dopamine should produce a smaller fluorescent change in hDAT-expressing GRAB_DA1M_ sniffer cells than in GRAB_DA1M_ sniffer that were mock transfected with an empty vector. To test this, we plated GRAB_DA1M_ sniffer cells that were transiently transfected with hDAT or empty vector as control and recorded the GRAB_DA1M_ fluorescence in a plate reader during stimulation with 1µM DA (**Figure 2A**). As expected, we observed a markedly lower fluorescent signal in cells expressing hDAT than in the mock transfected cells, reflecting hDAT-dependent DA uptake. To ensure that the reduced signal was truly mediated by hDAT and not an artifact of decreased surface expression levels of the sensor, the experiment was repeated in the absence and presence of the DAT blockers nomifensine and cocaine (**Figure 2B**). Indeed, pre-incubation with either blocker dose-dependently reversed the decreased fluorescent change upon addition of DA with a pIC_50_ of 6.09 ± 0.07 (820 nM) for cocaine and 6.77 ± 0.06 (170 nM) for nomifensine, which is comparable to earlier findings ^18^. Collectively these data demonstrate that the sniffer cells can be used as a radiotracer-free alternative for measuring DAT-dependent DA uptake and for screening of DAT blockers to derive indirect measures of apparent affinities.

**Figure 2.**
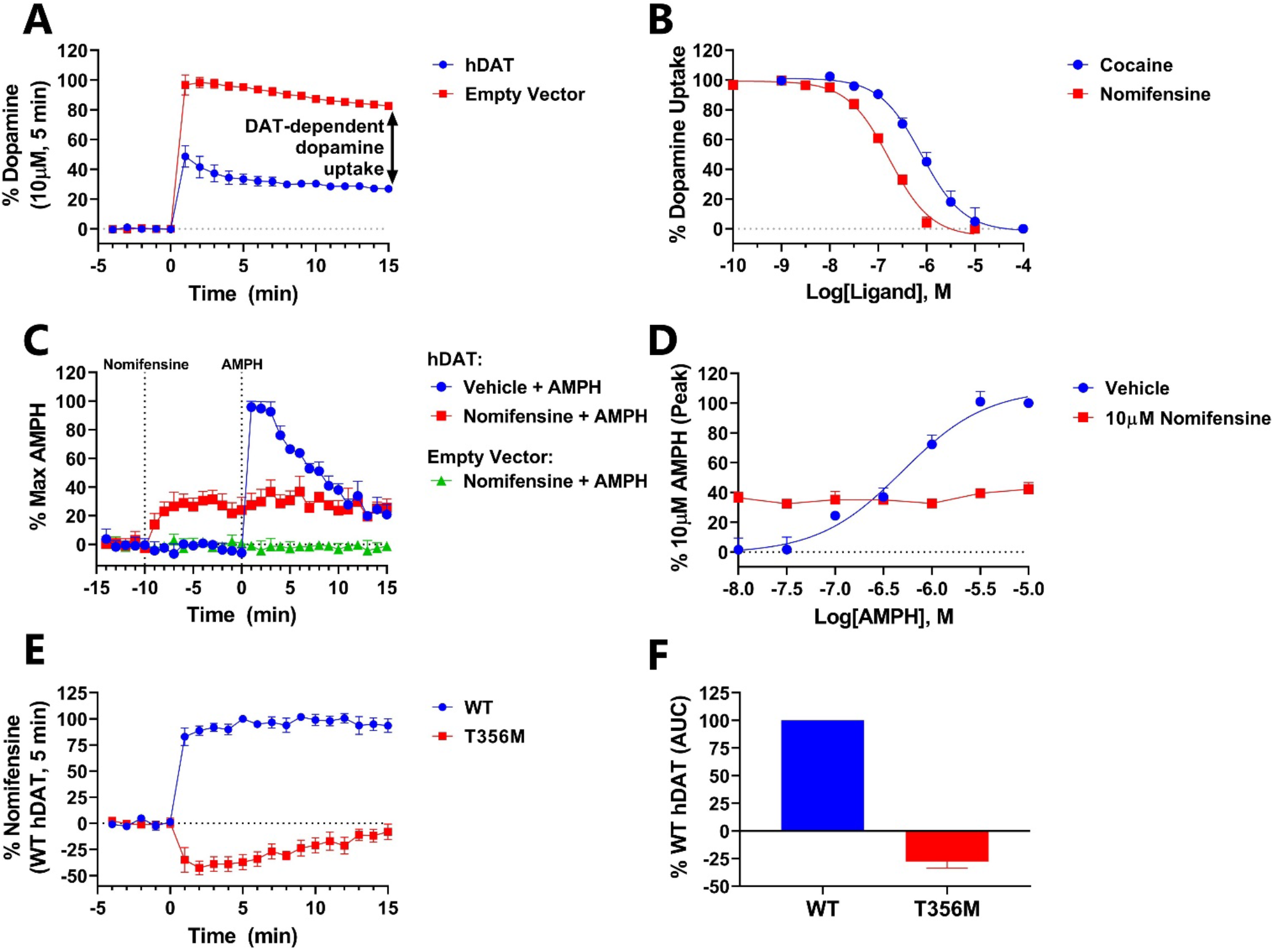
Detection of DAT-Mediated DA Uptake and Efflux Using Sniffer Cells. **A)** Measurement of DA uptake using DA sniffer cells. GRAB_DA1M_ sniffer cells, transfected with hDAT or an empty pcDNA3.1 expression vector, were stimulated with 1µM DA for 15 min. The decreased fluorescence upon addition of DA to cells transfected with hDAT versus an empty vector is indicative of DAT-mediated DA uptake into the cells. **B)** Pre-incubation of hDAT-transfected GRAB_DA1M_ sniffer cells with cocaine or nomifensine dose-dependently decreased the hDAT-mediated DA uptake. **C, D)** Measurement of AMPH-induced DA efflux via DAT using DA sniffer cells. GRAB_DA1M_ sniffer cells transfected with hDAT or an empty pcDNA3.1 expression vector were loaded with 300nM DA and washed subsequently. After reaching equilibrium, the baseline was recorded and the sniffer cells were incubated with 10µM nomifensine or vehicle for 10 min followed by a 15 min stimulation with AMPH (**C**, 10µM). AMPH caused a dose-dependent increase in fluorescence in hDAT-transfected cells, which was absent when the cells were preincubated with 10µM nomifensine (**D**). **E, F)** Recordings of anomalous DA efflux. GRAB_DA1M_ sniffer cells were transfected with WT hDAT or hDAT-T356M. After loading the cells with 10µM DA and subsequently washing away extracellular DA, the cells were stimulated with 10µM nomifensine. All values are expressed as mean ± SEM from three independent experiments conducted on a fluorescence plate reader. The data is expressed as normalized Δ(F/F_0_).

While DAT normally functions through inward transport of DA, studies on the mechanisms of psychostimulants have revealed that DAT can mediate reverse transport of dopamine as well, and that this efflux is essential for the action of AMPH ^19, 20^. Additionally, it has been shown that certain disease-associated mutations in DAT can cause anomalous constitutive DA efflux, which compromises the ability to accumulate DA ^21, 22, 23, 24^. We examinedwhether the sniffer cells could be used as an approach for detecting hDAT-mediated DA efflux.

First, we determined if we could observe AMPH-induced DA efflux in hDAT-transfected GRAB_DA1M_ sniffer cells in a high-throughput plate reader format. For this, sniffer cells were transfected with either hDAT or an empty vector and loaded with 300 nM DA for 15 min, after which extracellular DA was removed. We then added either vehicle or 10 µM nomifensine (10 min) before stimulating the cells with increasing concentrations of AMPH (10 nM – 10 µM). As seen in Figure 2, AMPH elicited a rapid dose-dependent increase in extracellular DA levels, measured as an increase in GRAB_DA1M_ fluorescence in hDAT-transfected cells, which was blocked by pre-incubation with nomifensine (**Figure 2C-D**). Moreover, from the dose-response curves we derived a pEC_50_ value of 6.32 ± 0.09 (476 nM) for AMPH (**Figure 2D**), which is comparable to what has previously been reported for hDAT-expressing heterologous cells ^25, 26^. It should be noted that the addition of nomifensine to hDAT-expressing GRAB_DA1M_ sniffer cells also increased the extracellular DA concentration, although markedly less than AMPH (**Figure 2C**). This increase presumably reflects DAT-independent DA leakage from the cells that can no longer be transported back into the cells by hDAT when blocked by nomifensine ^27^. Importantly, nomifensine and AMPH had no effect in GRAB_DA1M_ sniffer cells co-transfected with an empty vector, confirming that the observed effects are mediated through hDAT and not due to a direct interaction between the drugs and the sensor.

We then determined whether the sniffer cells could be used to validate an aberrant molecular phenotype, which has been described for an autism-associated *de novo* variant, hDAT-T356M. The hDAT-T356M variant imposes conformational changes to DAT that causes a leak of DA through the transporter (an anomalous DA efflux), which can be inhibited by DAT blockers ^21, 28^. To study this phenomenon with the sniffer cells, GRAB_DA1M_ sniffer cells were transiently transfected with either WT hDAT or hDAT-T356M, and loaded with a high DA concentration (10 µM) for 15 min. The cells were then washed and equilibrated for 20 min after which they were treated with nomifensine (10 µM) to block both reuptake and anomalous DA efflux. As expected nomifensine caused an increase in extracellular DA in WT DAT transfected cells as it blocks the reuptake of DAT-independent DA leakage. In contrast, stimulation of T356M-transfected sniffer cells with nomifensine produced a remarkable decrease in extracellular DA, consistent with blockage of constitutive DA efflux via hDAT-T356M (**Figure. 2E-F**). The phenomenon of anomalous DA efflux has been proposed to be a common mechanism through which missense mutations in DAT may exert disturbances in DA neurotransmission that are of pathophysiological relevance ^24^. So far, investigations of DAT-mediated DA efflux have relied on amperometric recording and superfusion assays with ^3^H-MPP^+ 21, 22, 23, 29^. Our data show that sniffer cells can be applied as an alternative, less labor-intensive strategy for identifying disturbances in DAT efflux properties and for studying the molecular and cellular consequences.

### Detection of DA Release from Cultured Dopaminergic Neurons and Striatal Slices, and Quantification of DA Content in Brain Tissue

Having established potential applications of the sniffer cells in heterologous cell assays, we next sought to explore if the sniffer cells could be applied to visualize release of endogenous DA from cultured dopaminergic neurons, and for *ex* vivo measurements of DA release from striatal slices and quantifications of DA tissue content in striatal homogenates.

To visualize DA release from cultured dopaminergic neurons, GRAB_DA2M_ sniffer cells were seeded on top of cultured neurons 24 h prior to experiments (**Figure 3A**). Wide-field fluorescence imaging was then conducted in aCSF (artificial cerebrospinal fluid) under constant slow perfusion. We stimulated the neurons with either KCl (**Figure 3B, Supplementary Video 1**) or electrical field stimulation (**Figure 3C**) to induce DA release and recorded the change in fluorescence of the GRAB_DA2M_ sniffer cells. Upon stimulation with either KCl or electrical field stimulation, an instantaneous increase in extracellular DA levels was detected. Termination of the electrical field stimulation reverted the fluorescent change back to baseline (**Figure 3C**). The decrease in fluorescent signal is presumably a combination of the speed of the perfusion system and the reuptake kinetics of DA back into the neurons.

**Figure 3.**
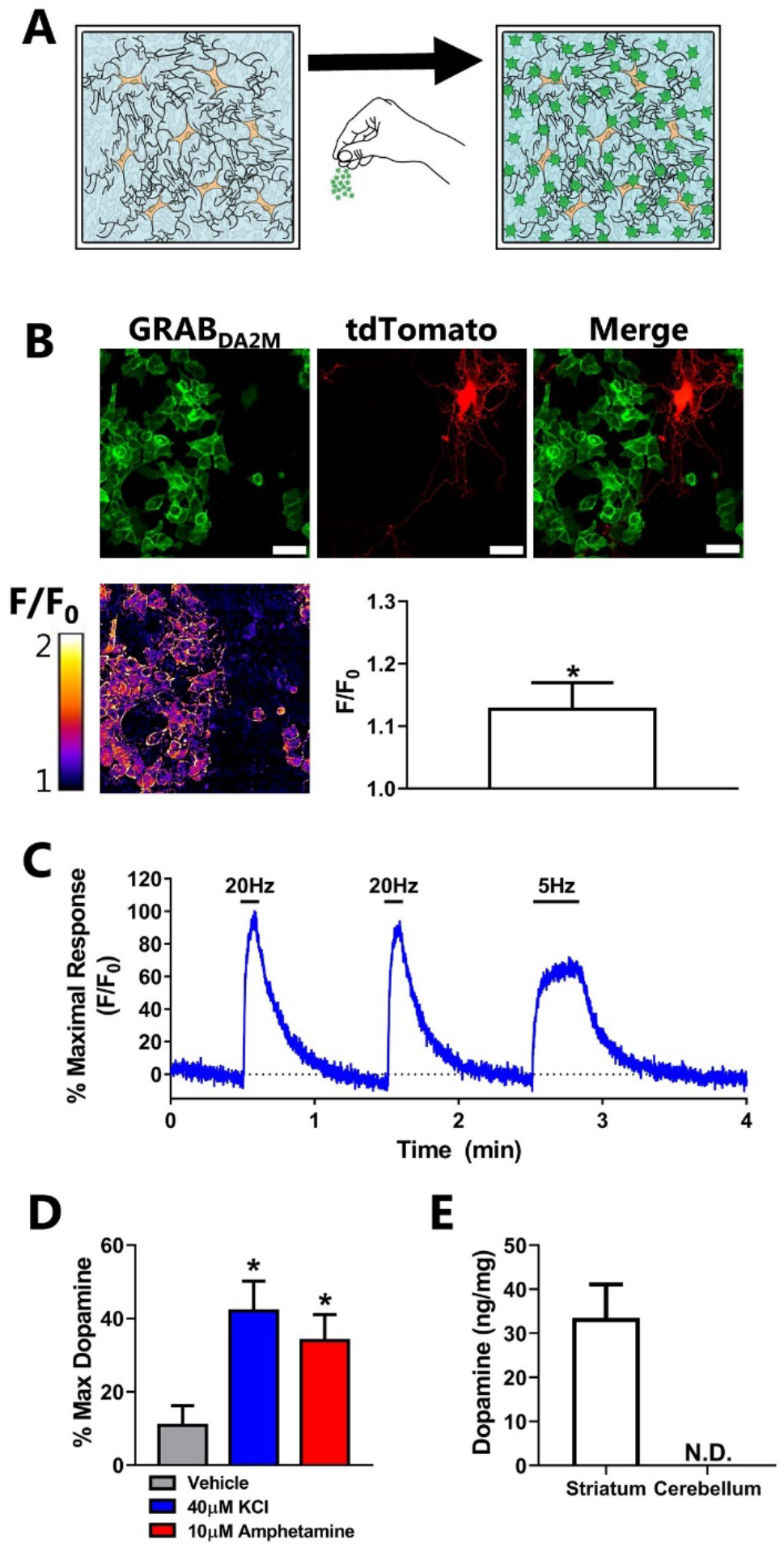
Detection of DA Release From Cultured Dopaminergic Neurons and Striatal Slices, and Quantification of DA Content in Brain Tissue. **A)** In order to detect DA release from cultured dopaminergic neurons, GRAB_DA2M_ sniffer cells were plated on top of the neurons 24 h prior to experiments. **B**) DA release from tdTomato-expressing dopaminergic neurons was induced by stimulation with 90mM KCl. The top row shows representative images of the GRAB_DA2M_ sniffer cells (green) and the tdTomato-expressing dopaminergic neuron (red). The bottom row shows a representative pseudo colored representations of the fold change above baseline in response to 90mM KCl, and also shows the fluorescent change detected in 5 recordings from 3 independent neuronal cultures. *P<0.05, one-sample t test. **C**) DA release evoked by electric field stimulation of 100 depolarizing monopolar pulses at a frequency of 20 or 5 Hz. DA release was detected as an increase in fluorescence from co-cultured GRAB_DA2M_ sniffer cells. The trace is representative of three independent experiments, and the data is expressed as the % F/F_0_ of the maximal response induced by the electrical field stimulation. **D)** Mouse striatal slices, incubated for 5 minutes (37°C) with vehicle, 40µM KCl, or 10µM AMPH in a small volume of aCSF. DA release was determined by transferring the aCSF incubation buffer to a 96-well plate seeded with GRAB_DA1M_ sniffer cells. DA content in the vehicle-, KCl-, and AMPH-treated samples was detected on a fluorescence plate reader, and the values are expressed as Δ(F/F_0_) normalized to the maximal Δ(F/F_0_) elicited by saturating [DA]. The values are the mean ± SEM from striatal slices of nine mice. *P<0.01, One-Way ANOVA with Dunnett post hoc test **E)** Total DA content was determined in striatum and cerebellum of mice utilizing the GRAB_DA1M_ sniffer cells. The tissue was homogenized, sonicated, and freeze/thawed in a hypotonic buffer to ensure that all DA was released from the tissue. Following centrifugation, the DA and protein concentrations were determined in the supernatant with a fluorescence plate reader allowing conversion to ng DA per mg protein. The DA levels in cerebellum were too low to be detectable by the GRAB_DA1M_ sniffer cells. The values are the mean ± SEM from tissue of eight mice.

Next, we tested if we could employ the sniffer cells to detect DA release from acute striatal slices. Striatal slices were submerged in aCSF and incubated for 5 min with either KCl or AMPH to induce DA release, or with vehicle. The buffer was then collected and added to GRAB_DA1M_ sniffer cells already seeded into 96 well plates and the change in fluorescence was detected with a fluorescence plate reader (**Figure 3D**). As expected, a greater increase in fluorescence was observed upon addition of media collected from KCl- and AMPH-treated slices than from vehicle-treated samples (N=9, P<0.01, **Figure 3D**).

Finally, we wanted to evaluate whether the sniffer cells could be used to determine DA content in mouse striatal tissue as an alternative approach to High-Performance Liquid Chromatography (HPLC) analysis. To do so, we prepared tissue homogenates from striatum in the absence of chemicals that could potentially affect the sniffer cells. As a negative control, we also prepared homogenates from cerebellum. In order to determine the DA concentration, the homogenates were added to GRAB_DA1M_ sniffer cells together with a DA standard curve (for interpolation), and the change in fluorescence was detected with a fluorescence plate reader. The average DA level detected in the striatal samples was 33.5 ± 7.6 ng/mg protein (mean ± S.E.M.; eight mice) while the DA level in the cerebellum was below the detection limit of the GRAB_DA1M_ sniffer cells (**Figure 3E**). The amount of DA detected in the striatal samples was within range of what has previously been reported by studies that utilized HPLC ^30, 31, 32, 33^. Thus, the sniffer cells provide a novel, alternative approach for determining total tissue DA content.

Taken together, the data presented show that sniffer cells can be used as an alternative method to visualize and measure DA release from dopaminergic neurons and striatal slices (e,g, to study the effect of drugs or genetic manipulations on DA release from dopaminergic release sites). Furthermore, the cells can also be utilized as a readily approachable way to determine total tissue DA content.

## Conclusions

Our study presents a framework for use of cells expressing genetically encoded fluorescence sensors as a virus- and radiotracer-free, inexpensive, and scalable approach for multimodal *in vitro* and *ex vivo* measurements of neurotransmitter levels. The sniffer cell strategy presented here should be a generalizable and easily applicable framework for other genetically encoded single-protein fluorescent sensors.

## Materials and Method

### Cloning

GRAB_DA1H_, GRAB_DA1M_ and GRAB_DA2M_ were subcloned into the pcDNA5/FRT/TO vector (Invitrogen) by Sequence and Ligation Independent Cloning (SLIC) using the procedure previously described^34^. The vector was linearized by cutting with EcoRV restriction enzyme, and inserts with the GRAB_DA_ sensors were generated by PCR with SLIC *Fwd* (5′-TGGAATTCTGCAGATATGGAGACAGACACACTC-3’) and *Rev* (5′-GCCACTGTGCTGGATTCAGCAGTGGAGGATCTT-3’) primers. dLight1.1 and dLight1.2 were both subcloned from their parent vector into the pcDNA5/FRT/TO vector using the HindIII and NotI restriction sites present in all three vectors. To generate dLight1.3a, we performed site-directed mutagenesis on dLight1.1 in pcDNA5/FRT/TO vector using the overlapping *Fwd* (5′-ACAGGATTGCTCAGAAACAGCTGAGCTCACTCATT-3’) and *Rev* (5′-AATGAGTGAGCTCAGCTGTTTCTGAGCAATCCTGT-3’) primers carrying the desired insert mutation (p.K247_L248insQ)^35^. All constructs were subsequently sequence verified to confirm correct insertion.

### Generation of Sniffer Cell Lines

Parental Flp-In T-REx 293 cells were grown in DMEM supplemented with 10% (v/v) fetal bovine serum (FBS; Invitrogen), 100 µg/mL zeocin (ThermoFisher Scientific), and 15 µg/mL blasticidin (ThermoFisher Scientific). To generate sniffer cell lines, the parental cells were grown in T150 flasks (Corning) until 70% confluency. The media was then changed to DMEM supplemented with 10% (v/v) FBS, and the cells were transfected with 0.6 µg DA sensor in a pcDNA5/FRT/TO vector and 5.4 µg pOG44 with 18 µL Lipofectamine 2000 (Invitrogen) according to the manufacturer’s protocol. 48 h after transfection, the cells were split 1:3 into a new T150 flask (Corning). After adherence of the cells, the media was changed to DMEM supplemented with 10% (v/v) FBS, 200 µg/mL Hygromycin B (Sigma), and 15 µg/mL blasticidin. The media was changed twice a week until colonies that stably express the sensor were obtained. Expression of the sensors was induced 24-48 h prior to experiments with 1 µg/mL tetracycline (Sigma).

### Culturing of Midbrain Dopaminergic Neurons

Cultures of midbrain dopaminergic neurons on top of cortical astrocytes were made from P1-P2 Wistar rats or DAT-IRES-Cre mice (Jackson Laboratory) as previous described ^36^. They were plated in 6-well plates on poly-D-lysine coated coverslips (Ø = 25 mm) using a modified serum-free medium (Neurobasal A (10888022, Gibco) with 1% GlutaMAX (35050061, Gibco), 2% B-27 plus (A3582801, Gibco), 200 µM ascorbic acid, 500 µM kynurenic acid and 0.1% Pen-Strep solution (P0781, Sigma). Two hours after plating neurons, rat glial cell derived neurotrophic factor (SRP3239, Sigma) was added for a final concentration of 10 ng/mL. The cultures were used for experiments 14-21 days after the neurons was plated out. The neurons obtained from DAT-IRES-Cre mice were transduced 5 days post dissection with pAAV-FLEX-tdTomato (Addgene).

### Imaging Experiments and Analysis

Most imaging was performed using an ECLIPSE Ti-E epifluorescence/TIRF microscope (NIKON, Japan) with a 488nm laser (coherent, California, USA) and an S Plan Fluor ELWD 20X/0.45 ADM microscope objective (NIKON, Japan). A 525/40 nm bandpass filter was used for the emission light, which was then recorded using an iXon3 897 Electron Multiplying CCD camera (Andor, United Kingdom). However, the imaging of the neurons obtained from DAT-IRES-Cre mice was conducted on a Nikon Eclipse FN1 upright microscope (Nikon, Japan).

For dose-response and max-response images of the individual DA sniffer cell lines, we seeded the cells out in 8-well Lab-Tek™ II Chambered Coverglass (Nunc) at a density of 25,000 cells per well and added tetracycline. The following day, the cells were washed twice with PBS and recorded with stepwise increasing [DA] for 5 minutes at each condition at a frame rate of 0.2Hz.

To record DA from neuron cultures, GRAB_DA2M_ sniffer cells were seeded out at a density of 200,000 cells per well on top of primary DAergic neuron cultures accompanied with tetracycline 1-2 days before the experiment. On the microscope, the neurons were mounted in a RC-21BRFS Field Stimulation Chamber (Warner Instruments, USA) and the neurons were continuous perfused with aCSF (in mM: NaCl, 120; KCl, 5; glucose, 30; MgCl_2_, 2; CaCl_2_, 2; HEPES, 25; pH 7.40) at 37°C. A Master-8 Pulse Generator (A.M.P.I., Israel) and an ISO-Flex Stimulus Isolator (A.M.P.I, Israel) were used to evoke DA release by passing 1 ms monopolar current pulses through the stimulation chamber electrodes to yield an electric field strength of ∼40 V/cm. Images were recorded at a frame rate of 12.5 Hz

All microscopy images were analyzed using the ImageJ-based Fiji software^37^. For calculating max-responses, we generated an averaged image of five consecutive images recorded for both conditions (before and one after adding 10 µM DA). Then, the pixel intensities over the same cellular cross-sections for each condition was quantified and single peak values (arising at cell membrane sections) were selected to calculate F/F_0_. For each N, we pseudo-randomly selected a population of cells with a low and a high baseline fluorescence and included two intensity peaks for each population. Thus, each N represents the mean F/F_0_ at four intensity peaks. The images of sniffer cells on neuron cultures were quantified by drawing a region of interest around a population of sniffer cells and measuring its intensity as a function of time using the build-in “Measure Stack” ImageJ macro.

### Fluorescence Plate Reader Experiments and Analysis

Sniffer cells were plated at a density of 30,000 cells/well into poly-L-ornithine-coated white CulturPlate-96 plates (PerkinElmer) and induced with tetracycline 48 h prior to experiments. In experiments with hDAT-transfected sniffer cells, the cells were simultaneously transfected with 50 ng hDAT and 150 nL Lipofectamine 2000 (Invitrogen) per well according to the manufacturer’s protocol. Fluorescence was measured (485/520) with a POLARstar OMEGA plate reader (Biotek), and all experiments were conducted at 37 °C.

For characterization of the sniffer cell lines, the cells were washed with aCSF and incubated in aCSF for 15 min at 37 °C in the absence or presence of the DA receptor antagonists SCH23390 or haloperidol. Upon measuring the baseline fluorescence intensity, the cells were incubated with vehicle or increasing DA concentrations, and the fluorescence intensity was measured after 5 min. To detect the rapid increase or decrease of fluorescence intensity upon the addition of DA or antagonist, the drugs were injected into the well by the POLARstar OMEGA plate reader (Biotek).

To detect uptake of DA via hDAT into the sniffer cells, hDAT- or pcDNA3.1-transfected GRAB_DA1M_ cells were incubated for 15 min at 37 °C with aCSF with or without cocaine or nomifensine. Afterwards, the cells were stimulated with 1 µM DA and the fluorescence intensity was measured for 15 min. The cells were also stimulated with 10 µM DA to use for data normalization.

AMPH-induced hDAT-mediated DA efflux experiments were performed by loading hDAT-transfected GRAB_DA1M_ cells with 300 nM DA for 15 min at 37 °C. The cells were then washed 3 times with aCSF for 3 min each. After a 20 min incubation at 37 °C the cells were stimulated with vehicle or nomifensine for 10 min followed by the addition of AMPH. The fluorescence intensity was measured every minute throughout the experiment.

Experiments that allow the detection of constitutive hDAT-mediated DA efflux were performed in a similar manner as the AMPH-induced DA efflux experiments. However, the cells were loaded with 10 µM DA rather than 300 nM DA, and the cells were only stimulated with nomifensine and not with AMPH.

To analyze the data, the fluorescence intensity was divided by the mean baseline fluorescence intensity to obtain F/F_0_. Subsequently, the average F/F_0_ of control wells that solely received vehicle was subtracted to gain Δ(F/F_0_). These values were then normalized as indicated. All data was analyzed using GraphPad Prism 9.

### Detection of DA Release from Acute Mouse Striatal Slices

Acute brain slices were acquired from adult mice (14 ± 2 weeks). The animals were anesthetized with isoflurane and the brains were quickly harvested into ice-cold aCSF. Coronal striatal brain slices of 300 µm thickness were prepared on a LeicaVT1200 vibrating blade microtome. Slices were then transferred to oxygenated aCSF at room temperature and allowed to recover for at least 1h before the experiment. Subsequently, the slices were transferred to 2 mL Eppendorf tubes containing 500 uL pre-warmed (37 °C) aCSF in the presence or absence of 40 mM KCl or 10 µM AMPH. After 5 min incubation at 37 °C, the aCSF was collected to determine whether DA release had occurred.

To detect whether stimulation with KCl or AMPH induced DA release from acute striatal slices, GRAB_DA1M_ sniffer cells were utilized. The cells were washed with 200 µL aCSF and incubated for 15 min at 37 °C with 100 µL aCSF. After a baseline read, 100 µL aCSF collected from the slices was added to the cells, and the response was measured after 5 min on the POLARstar OMEGA plate reader. To normalize the data, cells were also stimulated with aCSF and 10 mM DA.

### Determination of DA Content in Mouse Striatum and Cerebellum

In order to extract DA from striatal tissue samples, we dissected striatal tissue samples from coronal slices using a brain matrix and a puncher. We also cut tissue samples from cerebellar slices to use as a negative control. The tissue was collected in hypotonic buffer (25 mM HEPES, pH 7.40 with KOH) containing 1 mM glutathione (Sigma) to prevent oxidation of DA ^38^. To ensure extraction of all DA, the sample was homogenized using a syringe with a 27 G needle, sonicated and freeze/thawed 5 times by alternating between 37 °C and −80 °C. The sample was then centrifuged for 30 min at 4 °C at 16 *g*, and the supernatant was collected for DA and protein determination.

The protein concentration was determined with a standard BCA kit (Pierce). The DA concentration was determined utilizing GRAB_DA1M_ sniffer cells plated in a white CulturPlate-96 plate (PerkinElmer). The cells were washed with 200 µL aCSF, and then incubated for 15 min at 37 °C in 100 µL aCSF. After a baseline read, 100 µL supernatant was added to the cells, and the response was measured after 5 min on the POLARstar OMEGA plate reader. To determine the DA concentration, additional wells with cells were also incubated with a range of DA concentrations prepared in hypotonic buffer to allow interpolation.

## Acknowledgements

We thank Anette Dencker Kaas for excellent technical assistance. The work was supported by the Independent Research Fund Denmark – Medical Sciences (C.K.H: 6110-00292B, UG: 7016-00325B, and UG: 9039-00437B). The Lundbeck Foundation (FH: R303-2018-3540, and UG: R276-2018-792)

## Author Contributions

C.K.H, J.F.S, and F.H. were responsible for the overall experimental design. C.K.H, J.F.S, W.D.R, A.D, M.A.C.G, and F.H conducted the experiments. C.K.H and F.H. wrote the manuscript. C.K.H, U.G., and F.H. provided funding. Y.L and L.T. developed and provided the sensors. All authors have contributed to the editing and review of the final manuscript.

