## Supplementary Material for "Multimodal Detection of Dopamine by Sniffer Cells Expressing Genetically Encoded Fluorescence Sensors"

[1]Molecular Neuropharmacology and Genetics Laboratory, Department of Neuroscience, Faculty of Health and Medical Sciences, University of Copenhagen, Copenhagen, Denmark. [2]College of Medicine, University of Florida, Gainesville, FL 32611, USA. [3]State Key Laboratory of Membrane Biology, Peking University School of Life Sciences, 100871 Beijing, China, [4]PKU-IDG/McGovern Institute for Brain Research, 100871 Beijing, China, [5]Peking-Tsinghua Center for Life Sciences, 100871 Beijing, China. [6]Departments of Biochemistry and Molecular Medicine, School of Medicine, University of California, Davis, Davis, CA, USA

[†]Address author correspondence to: Freja Herborg, Department of Neuroscience, Maersk Tower 7.5, University of Copenhagen, Blegdamsvej 3B, DK-2200 N, Copenhagen, Denmark. Phone +4553609699;

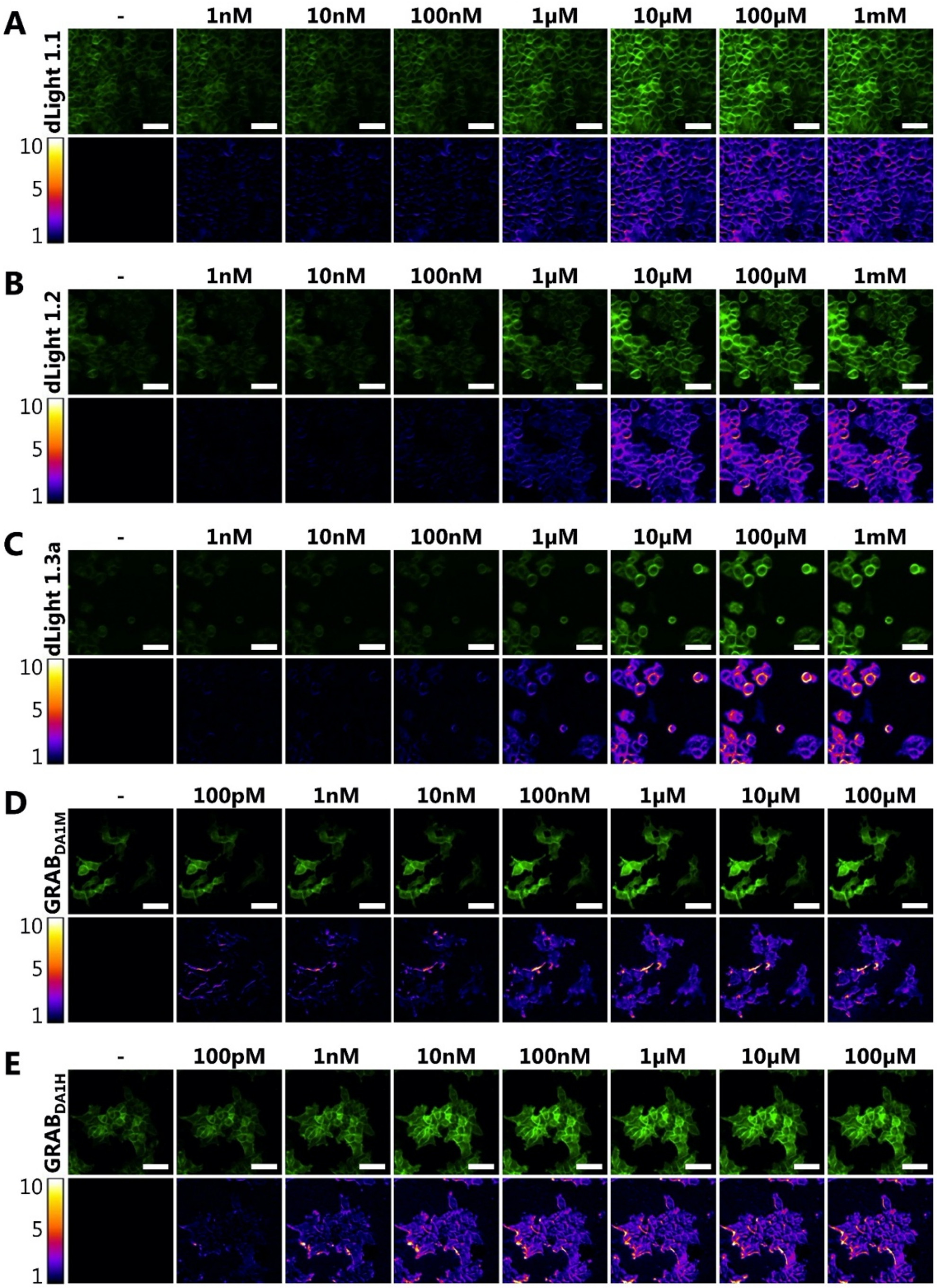


**Supplementary Figure 1 Characterization of the Different DA Sensor-Expressing Sniffer Cell Lines.** The dose-dependent increase in fluorescence upon addition of increasing DA concentrations to cells expressing the dLight1.1 (**A**), dLight1.2 (**B**), dLight1.3a (**C**), $GRAB_{DA1M}$ (**D**), and $GRAB_{DA1H}$ (**E**) sensors as detected by an epifluorescence microscope. Bottom panels show the change in fluorescence ($F/F_0$) upon addition of DA. Images shown are representatives from three independent experiments. Scale bars are 50μM.

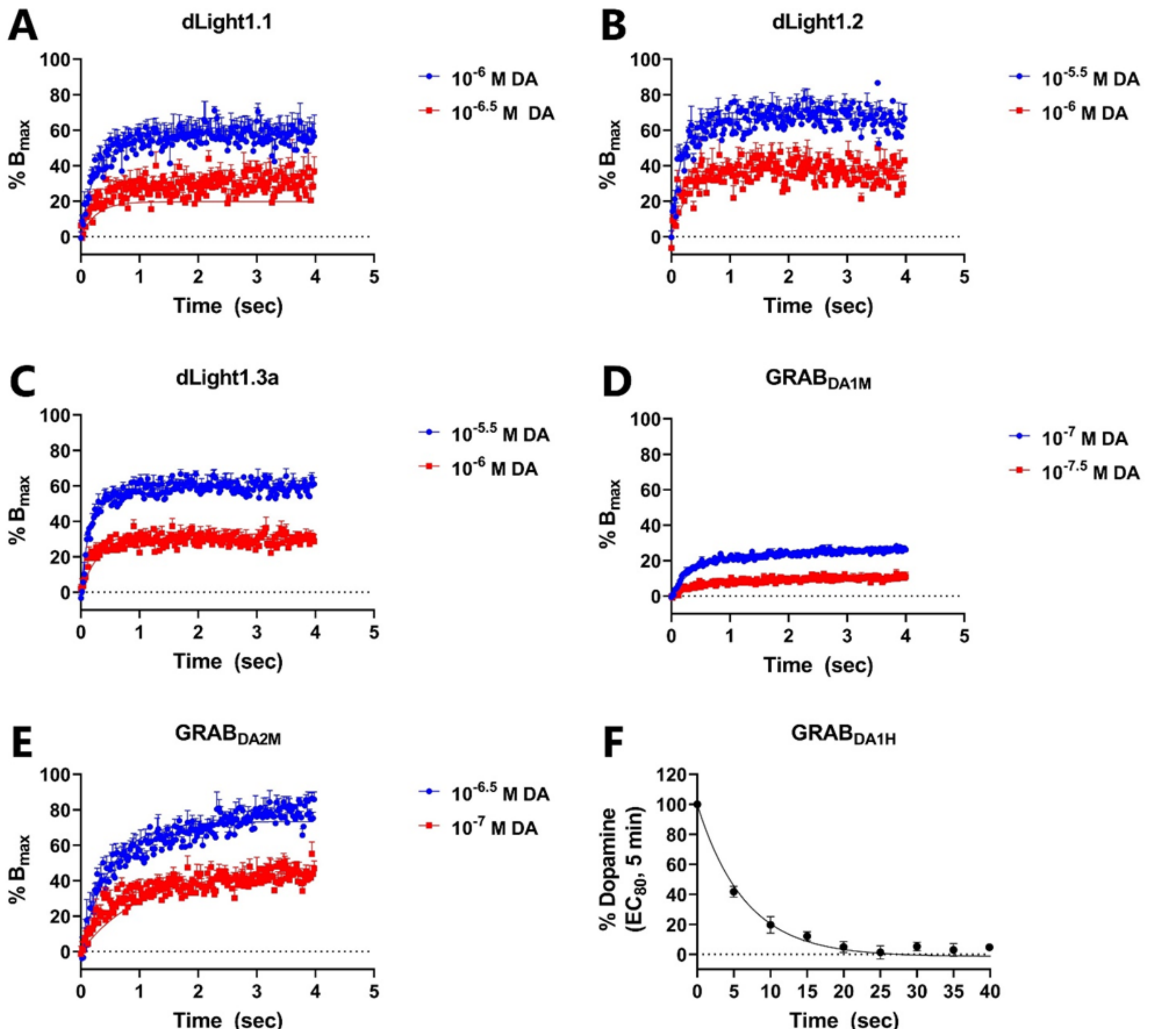


**Supplementary Figure 2 Determination of Kinetic Parameters of DA Sensors. A-E)** Sniffer cells were stimulated with two concentrations of DA and the change in fluorescence was determined over time. The data was then fitted with Graphpad Prism (association kinetics (two ligands concentrations)) to allow the determination of the on ($k_{on}$) and off ($k_{off}$) activation rates. **F)** $GRAB_{DA1H}$-expressing sniffer cells were stimulated for 5 minutes with an $EC_{80}$ concentration of DA followed by 10μM haloperidol (time-point 0) to determine the off-rate. The data was fitted with Graphpad Prism (Dissociation - One phase exponential decay). Values are expressed as mean ± SEM from three independent experiments conducted on a fluorescence plate reader.

**Supplementary Video 1** The video shows the fluorescence change of $GRAB_{DA2M}$ sniffer cells co-cultured with tdTomato-expressing mouse dopaminergic neurons upon stimulation with 90mM KCl.
